# Segregation between an ornamental and a disease driver gene provides insights into pigment cell regulation

**DOI:** 10.1101/2024.05.20.595041

**Authors:** Erika Soria, Qiusheng Lu, Will Boswell, Kang Du, Yanting Xing, Mikki Boswell, Korri S Weldon, Zhao Lai, Markita Savage, Manfred Schartl, Yuan Lu

## Abstract

Genetic interactions are adaptive within a species. Hybridization can disrupt such species-specific genetic interactions and creates novel interactions that alter the hybrid progeny overall fitness. Hybrid incompatibility, which refers to degenerative genetic interactions that decrease the overall hybrid survival, is one of the results from combining two diverged genomes in hybrids. The discovery of spontaneous lethal tumorigenesis and underlying genetic interactions in select hybrids between diverged *Xiphophorus* species showed that lethal pathological process can result from degenerative genetic interactions. Such genetic interactions leading to lethal phenotype are thought to shield gene flow between diverged species. However, hybrids between certain *Xiphophorus* species do not develop such tumors. Here we report the identification of a locus residing in the genome of one *Xiphophorus* species that represses an oncogene from a different species. Our finding provides insights into normal and pathological pigment cell development, regulation and molecular mechanism in hybrid incompatibility.

**Significance:** The Dobzhansky–Muller model states epistatic interactions occurred between genes in diverged species underlies hybrid incompatibility. There are a few vertebrate interspecies hybrid cases that support the Dobzhansky–Muller model. This study reports a fish hybrid system where incompatible genetic interactions are involved in neuronal regulation of pigment cell biology, and also identified a novel point of regulation for pigment cells.

## Introduction

The genome incompatibility hypothesis stated by Dobzhansky-Muller (DM) describes that negative genetic interactions in hybrids serve as the molecular mechanisms accounting for hybrid incompatibility (1–5). Almost a century ago, three independent studies found that interspecies hybrids between southern platyfish *Xiphophorus maculatus* (*X. maculatus*) and green swordtail *X. hellerii* develop spontaneous melanoma (6–8). This fish model represents one of the examples supporting the DM incompatibility hypothesis (9–11). The *X. maculatus* exhibits a nevus-like black pigmentation pattern (*Spotted dorsal*, or *Sd*), and a red pigmentation pattern (*Dorsal red*, or *Dr*) in the dorsal fin, while the *X. hellerii* exhibits neither trait. In hybrids between the *X. maculatus* and *X. hellerii*, the *Sd* and *Dr* pigmentation pattern becomes enhanced and expanded. The *Sd* pattern covers the entire dorsal fin and *Dr* pattern expands to the tail fin and posterior of the body from dorsal fin (12, 13). In the backcross hybrid using the *Sd* and *Dr* negative species (i.e., *X. hellerii*) as recurrent parental, 50% of the hybrids inherited recurrent parental species pigmentation pattern (i.e., no *Sd*, nor *Dr*), 25% of hybrids exhibit an F_1_-like *Sd* pattern, and 25% exhibit invasive and lethal nodular exophytic melanoma (12–14). The *Dr* pattern never progressed to a pathological condition.

It has been found that the *X. maculatus Sd*, the enhanced *Sd* pattern, and melanoma in the hybrids is driven by a mutant duplicate copy of Epidermal growth factor receptor (EGFR) that is termed *<u>X</u>iphophorus <u>M</u>elanoma <u>R</u>eceptor Tyrosine <u>K</u>inase* (*xmrk*) (15, 16). The expressivity of *Sd*, however, is regulated by copy number of a co-evolved *xmrk* regulator in *X. maculatus*. With two copies of the *xmrk* regulator, *Sd* is restricted to a nevus-like pigmentation dot, while losing the *xmrk* regulators through hybridization leads to pigment cell benign hyperplasia (losing one copy of the regulator), and melanoma (losing both copies of the regulator). The *xmrk* regulator has recently been mapped to chromosome 5 of *X. maculatus* genome (13).

The *Sd* and *Dr* loci in the *X. maculatus* offer a naturally occurred genetic system by which an oncogene activity can be assessed by phenotyping pigmentation patterns. The *Dr* is thought to be tightly linked to *Sd* as *Dr* co-segregates with *Sd* in the *X. maculatus-X. hellerii* hybrids. However, the *Dr* does not exhibit an *Sd-*like dosage effect from the *xmrk* regulator. The *Sd* is a melanophore phenotype, while the *Dr* is composed of xanthophores. The two cell types are in different differentiation trajectories of neural crest cell through *pax7a* and *pax7b* dependent fate determination (17). The *Dr* does not develop into tumor while *Sd* can progress into melanoma. This contrasts to the situation in the *mitf* promoter driven *xmrk* transgenic medaka, which develop several types of pigment cell tumors. In these fish the *xmrk* expression is driven by the *mitf* promoter that is active in xanthophores and melanophores. As a result, tumors formed in both cell lineages. Even so, the xanthophore tumor (i.e., xanthophoroma) only develops as epiphytic nodules without invasion. Xanthophoroma gene expression profile suggests it is less proliferative and less invasive than the melanoma counterpart (18, 19). The absence of the *xmrk* effect in xanthophores in *Xiphophorus* hybrids implies that despite *Sd* and *Dr* being linked yet are subjected to different regulation.

The genetics underpinning the *Dr* and *Sd* has not been resolved. Therefore, to study the discrepancy between *Sd* and *Dr*, we developed F_2_ intercross interspecies hybrid between *X. maculatus* and *X. couchianus*. *X. maculatus* and *X. couchianus* are distantly related species. They are historically classified to southern and northern platyfish based on their habitat and phylogeny based on partial genome sequences (20). Morphologically, the two species are distinguished by body shape and pigmentation patterns. Especially the *X. maculatus* has *Sd* and *Dr*, while *X. couchianus* doesn’t possess either phenotype (Fig. 1). Importantly, unlike the aforementioned *X. maculatus* and *X. hellerii* hybrid, *X. maculatus* and *X. couchianus* F_1_ hybrid only exhibit *Dr*, but not *Sd.* Therefore, it offers a genetic model system that can be used to study diverged regulation of *Dr* and *Sd.* This *Dr* and *Sd* expressivity divergence leads to the hypothesis that *X. couchianus* genome encodes a dominant suppressor of *xmrk-* driven *Sd*. Herein this study, we first tested whether *Sd* can express in F_2_ interspecies hybrid population where a hypothetical dominant *X. couchianus xmrk* suppressor is conditionally lost, and subsequently investigated genetic mechanism underpinning of the *Sd* and *Dr*.

**Figure 1.**
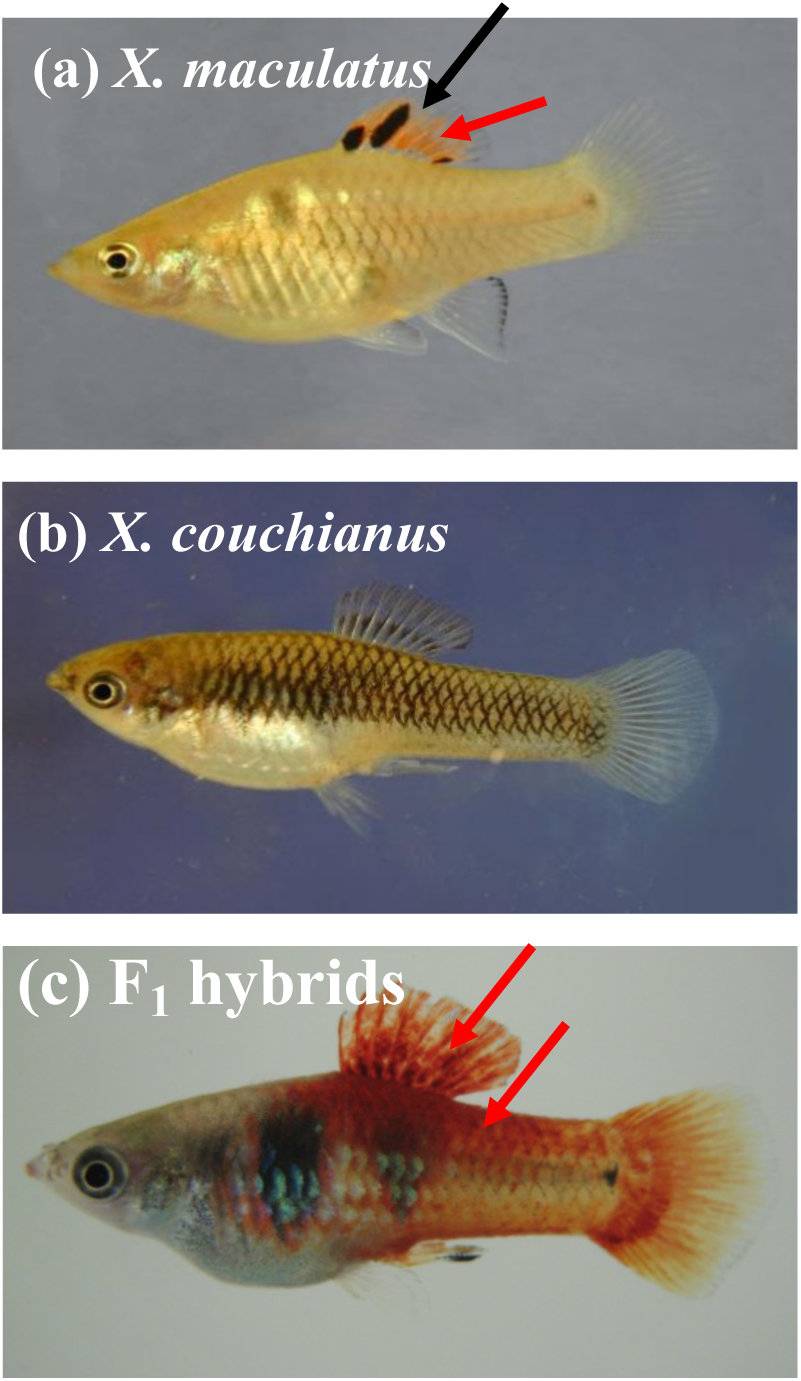
Pigmentation patterns of *X. maculatus*, *X. couchianus* and their hybrid. (a) The *X. maculatus* exhibit a large dorsal fin spotting pattern that is encompassed by macromelanophore (*Spot Dorsal,* or *Sd*, pointed by black arrow). The dorsal fin also exhibits xanthophore-driven red coloration (*Dorsal Red,* or *Dr*, pointed by red arrow). (b) The *X. couchianus* does not exhibit *Sd* or *Dr*, but only background color made by micromelanophore. (c) The F_1_ interspecies hybrids between the *X. maculatus* and *X. couchianus* display enhanced and expanded xanthophore pigmentation (pointed by red arrows), but not *Sd*. F_2_ images are included in Supplemental Data.

## Results

### Pigmentation pattern divergence in F_2_ interspecies hybrids

In contrast to the analogous cross between *X. maculatus* (*Sd +*, *Dr* +) and *X. hellerii* (*Sd -*, *Dr* -), *Dr* is enhanced in *X. maculatus* and *X. couchianus* (*Sd -*, *Dr* -) F1 interspecies hybrids (*mac-cou* hyb) while *Sd* is not expressed in neither the *mac-cou* F_1_ nor backcross hybrid using *X. couchianus* as recurrent parent (Supplemental Table S1, Supplemental Table S2). We hypothesized that the *X. couchianus* genome encodes a dominant locus suppressing the *Sd* but not *Dr.* Therefore, we created a *mac-cou* F_2_ interspecies hybrid population to assess *Sd* and *Dr* inheritance. We produced a total of 93 F_2_ interspecies hybrids (*mac-cou hyb*). A 9.5% of the F_2_ intercross hybrids exhibit *Sd*, compared to 64.8% for *Dr* (Supplemental Table S3). All hybrids exhibiting *Sd* have *Dr,* but not vice versa.

We compared the skin gene expression profiles of the *mac-cou hyb* exhibiting *Sd* to the hybrids exhibiting only *Dr* and identified differentially expressed genes between the two pigmentation patterns. There are 28 genes over-expressed in *Sd*, including, as expected, genes related to the macromelanophore, melanocyte and melanoma (*slc24a5*, *st3gal3*, *st6galnac3*, *dct*, *oca2*, *melanopsin*-*A*-*like*, *fam19a1*, *tyrp1*, *pmel*, and *emilin2*). An unannotated gene, LOC11161040 that is the homolog to long neurotoxin OH-57 is the only gene under-expressed in *Sd* hybrids (Table 1).

**Table 1:**
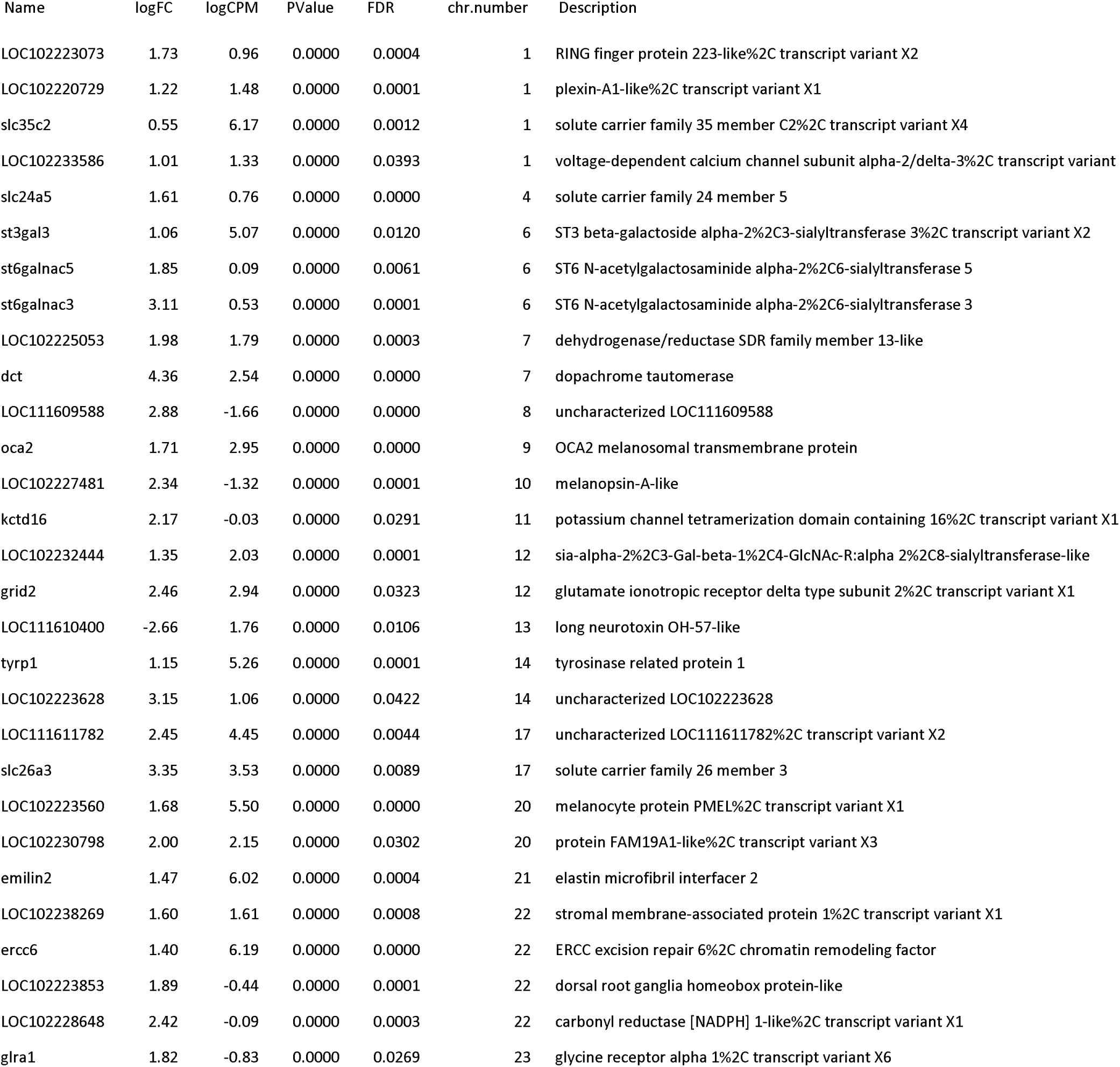
Differentially expressed genes between *Sd* and *Dr* hybrids skin.

### Segregation between two oncogene-driven pigmentation patterns in interspecies hybrids

We estimated inter-species variant sites’ genotype and genetically mapped *Dr* and *Sd*. We categorized the F_2_ population based on presence of *Dr* (*Dr+* and *Dr-*) or *Sd* (*Sd*+ and *Sd*-). Mapping of *Dr* showed one single peak associated with the trait on chromosome (Chr) 21 between 24.27 to 25.03 Mbp. This region contains 18 gene models, including the *xmrk* (Fig. 2a). All F_2_ hybrids that are either homozygous for the *X. maculatus* alleles of genes within the peak region, or heterozygous for the peak region exhibited the *Dr* (Fig. 2a; 100% penetrance).

**Figure 2.**
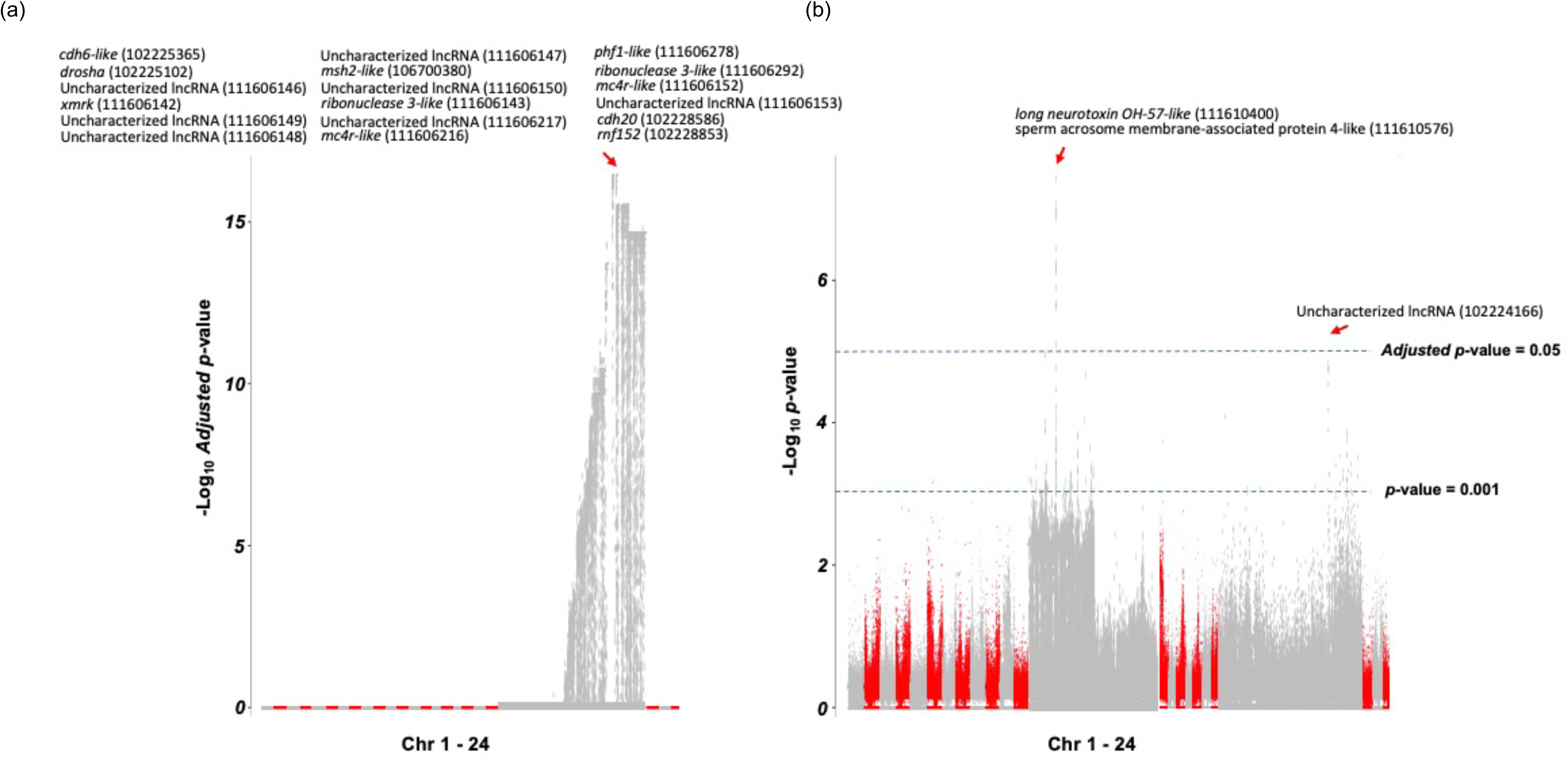
Genetic mapping identifies one locus underlying *Dr* and two loci underlying *Sd*. Interspecies hybrids were produced by mating male and female *X. maculatus-X. couchianus* F_1_ hybrids. (a) Genetic mapping showed one single peak on Chr 21 is associated to the *Dr.* In the Manhattan plot, X-axis represent all 24 chromosomes, with gray represented odd number chromosomes and red represented even number chromosome. Non-Chr 21 chromosomes were reduced in size as there is no statistically significant locus identified on them. Y-axis represent multiple test correction adjusted p-values that are transformed to - 1Log_10_ Adjusted p-value. The *Dr* is mapped to 24.27 to 25.03 Mbp on Chr 21. This region contains 18 genes. All *Sd+* F_2_ hybrids are either homozygous for the *X. maculatus* allele, or heterozygous for both parental alleles in the peak region. (b) The *Sd* is mapped to a region around 5.50 Mbp region on Chr 13 and 23.89 Mbp region on the Chr 21. The Chr 13 region contains 2 genes, and the Chr 21 region contains one gene. For the Chr 13 locus, 80% of the *Sd+* hybrids are homozygous for the *X. maculatus* allele, 20% are either homozygous for *X. couchianus* allele, or heterozygous for both parental alleles. For the Chr 21 locus, 20% of the *Sd* hybrid is homozygous for *X. maculatus* allele, and 80% of the *Sd* hybrid is heterozygous for both parental alleles.

In comparison, the *Sd* is associated with two loci, including a region located around 23.89 Mbp on Chr 21 that overlapped with one gene encoding an uncharacterized long noncoding RNA (Fig. 2b), and another loci around 5.50 Mbp region on Chr13 that located between *long neurotoxin OH-57 like* and *sperm acrosome membrane-associated protein 4-like*. For the Chr21 locus, 20% of the *Sd* hybrids are homozygous for the *X. maculatus* allele, and 80% of the *Sd* hybrids are heterozygous for both parental alleles (Supplemental Figure 2). In order to test if the Chr 21 loci underlying *Dr* and *Sd* are closely linked, we utilized a sub population of the F_2_ hybrids that exhibited *Dr*, and categorized this population as *Dr+, Sd-* and *Dr+, Sd+*. The Chr21 loci became insignificant in associating with *Sd*, indicating the loci underpinning the *Sd* and *Dr* are not resolved (Supplemental Figure 1).

The Chr 13 region associated with the *Sd* is preferentially, i.e., 80% penetrance, homozygous for *X. maculatus* allele (Fig. 3). All hybrid fish that exhibit *Sd* and are homozygous for *X. maculatus* allele showed extra macromelanophore pigmentation spots on trunk musculature (Supplement Data). The two fish samples that exhibit *Sd* but are heterozygous for Chr13 (Fish 23), or homozygous for *X. couchianus* Chr13 (Fish 48) showed atypical *Sd* pigmentation pattern (Fig. 3b,c). These melanophore clusters are smaller in size than those in *X. maculatus* parental. In addition, the macromelanophore distributed in a more dispersed pattern than the parental in which the macromelanophores aggregate to form a large spot (i.e., Fig. 3c).

**Figure 3.**
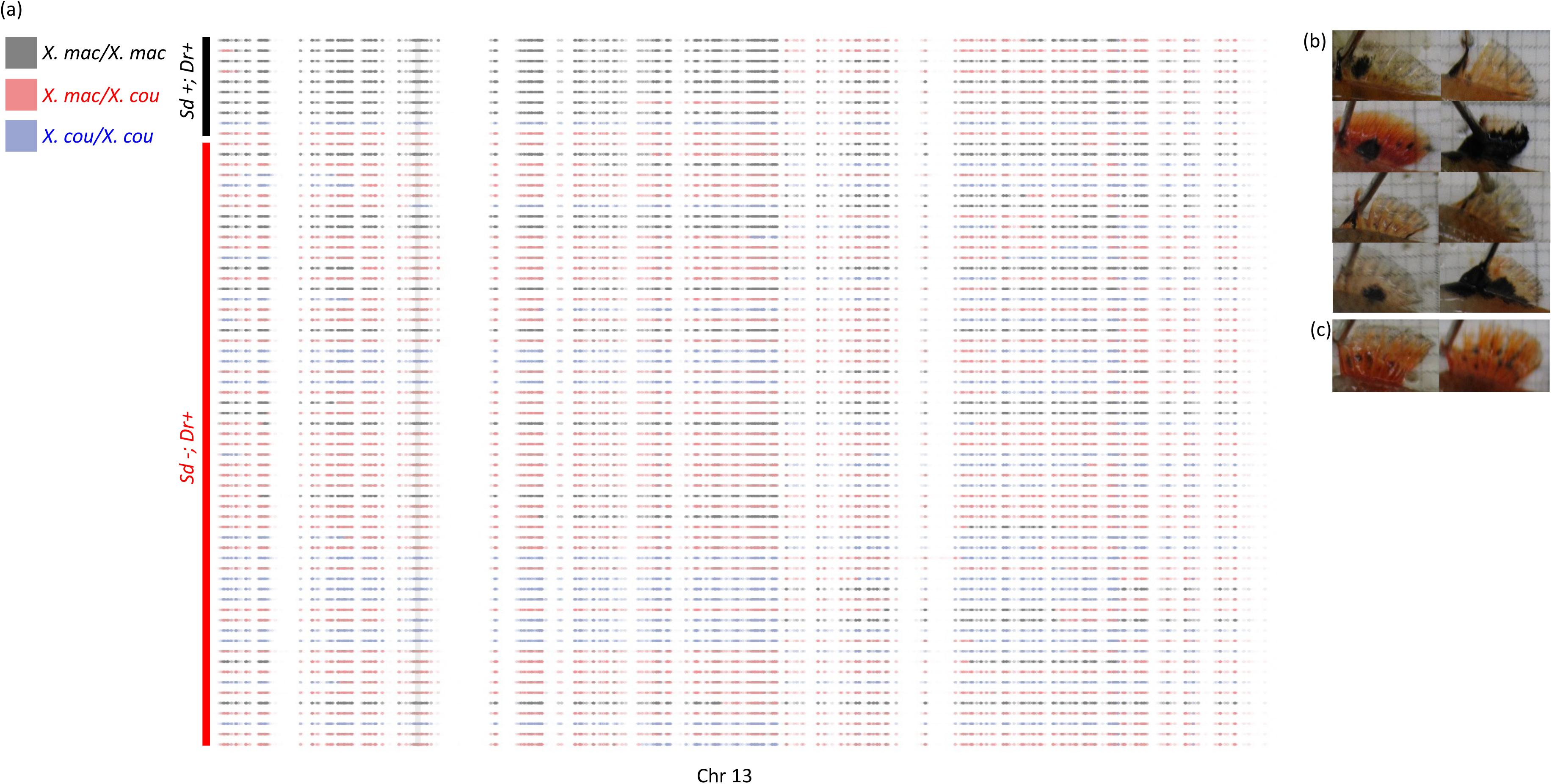
Chromosome recombination pattern does not support genetic segregation between *Sd* and *Dr*. (a) Haploid map for Chr 13 for *Dr+Sd*- and *Dr+Sd*+ hybrids were made by plotting genotype of inter-specific polymorphism genotypes along their chromosomal coordinates. Gray, red and blue dots are respectively polymorphic sites that are homozygous for *X. maculatus*, heterozygous for both parental alleles, and homozygous for the *X. couchianus* allele. Chromosomal recombination took place where bars of different colors meet. Vertical shaded gray line hall marks loci underlying *Sd*. (b) The melanophore patterns for 8 *Sd+* hybrids. (c) The melanophore patterns for 2 *Sd+* hybrids that exhibited irregular macromelanophore patterns. They are characterized by smaller melanophore aggregates and more dispersed micromelanophore pattern that is distributed between fin rays.

The definition of gene mapping relies on the total number of chromosome recombination. Therefore, we expected to observe that *Sd* hybrids would exhibit chromosomal recombination in loci that are adjacent to the Chr13 5.50 Mbp region. However, the chromosome recombination patterns showed by the haploid maps only support the shoulder region around the 5.50 Mbp peak (Fig. 3, Supplement Figure 2). Similarly, recombination did not occur between Chr21 locus underlying *Dr* and the locus associated with *Sd* (Supplement Figure 2).

### Allelic gene expression associated to *Dr* and *Sd*

For a gene to be functional, the minimum requirement is the expression of the allele underling the function. We hypothesized that the alleles associated to *Sd* are preferentially expressed by one allele in *Sd-* or *Sd+* hybrids. First of all, the *xmrk*, as a X*. maculatus* specific gene, showed the same level of *X. maculatus* allele specific expression in the hybrid expressing *Dr*, including the individuals that also display *Sd* (Supplement Figure 3). *Dr+ Sd-* hybrids exhibited bi-allelic expression for *cdh6, drosha, mc4r* and *cdh20*, while the hybrids without the *Dr* (*Dr- Sd-*) exhibit *X. couchianus* allele specific expression for *cdh6, drosha,* and *cdh20*, and no expression for *mc4r* (Fig. 4a,b). These findings support that the *Dr* and *Sd* are *X. maculatus* specific traits

**Figure 4.**
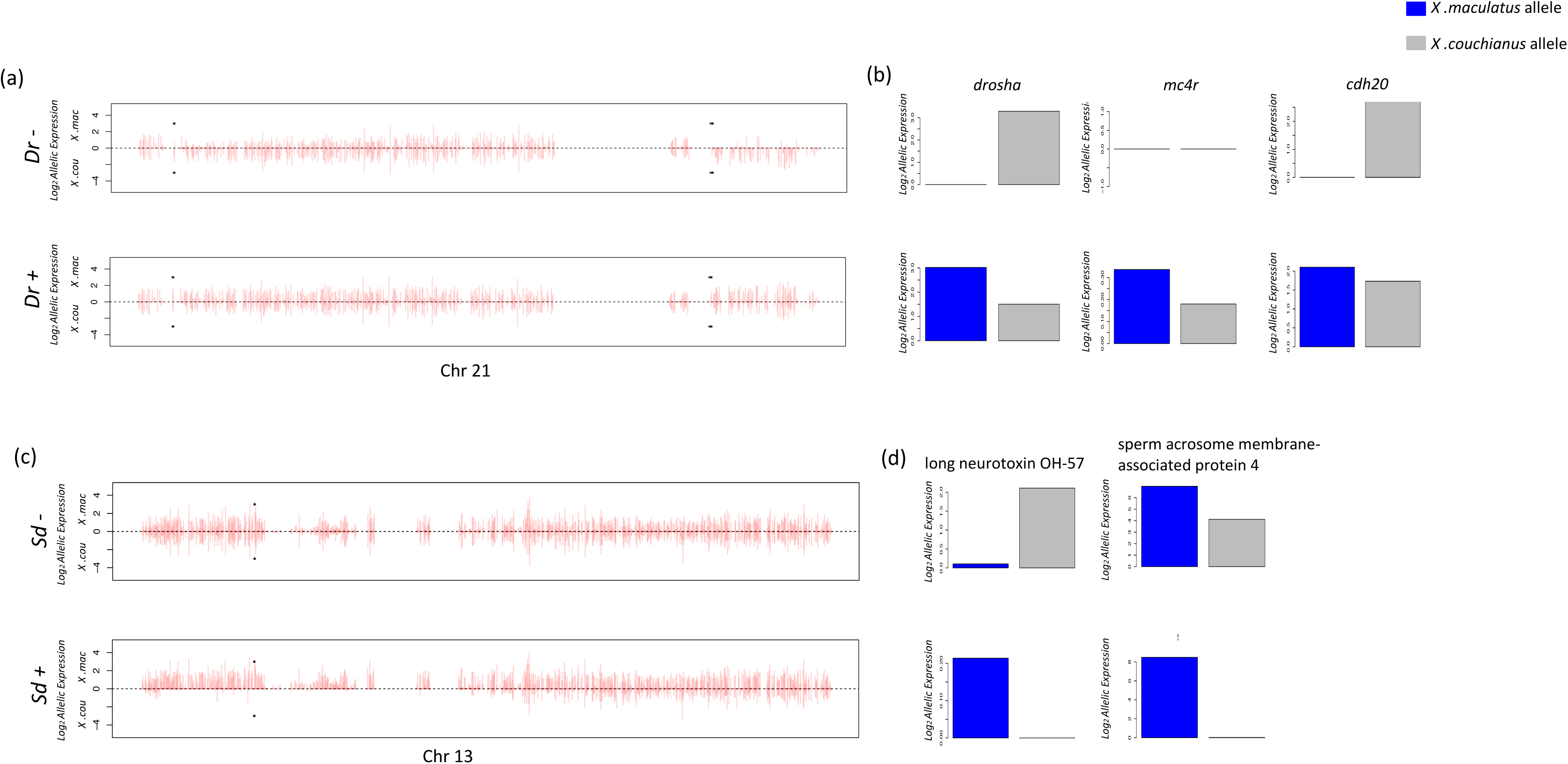
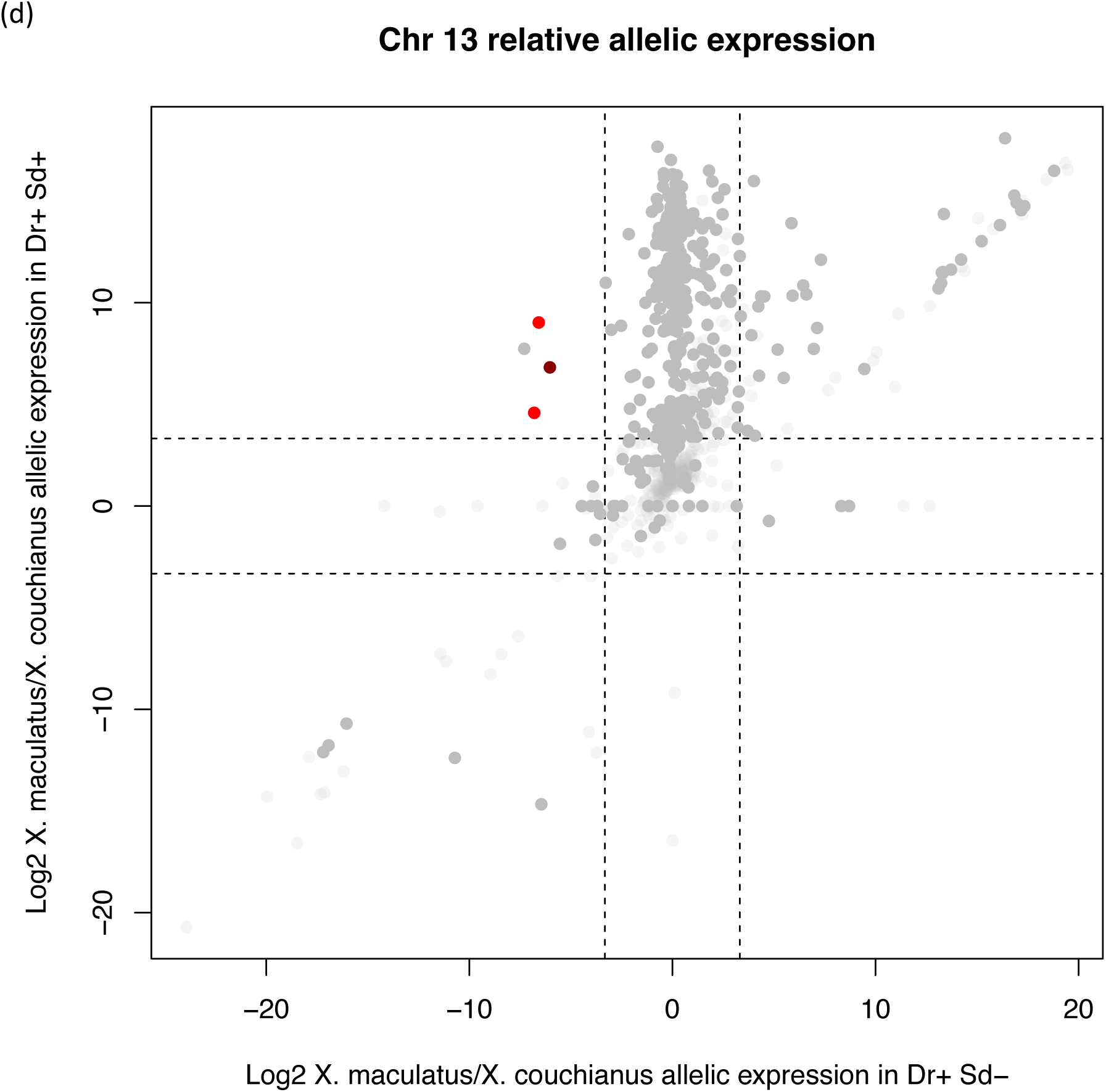
Allelic gene expression pattern hallmarks genes associated with *Dr* and *Sd*. (a) The upper panel and lower panel shows allelic expression in *Dr-* and *Dr+* hybrids respectively. Allelic expression values were normalized. Bars above the middle line represent the *X. maculatus* alleles, and bars under the middle line represent *X. couchianus*. alleles. Bars are plotted in the order of *X. maculatus* gene orders on Chr 21. Asterisks hallmark positions of candidate gene models. Because the Chr 21 assembly has a double-inversion assembly error, the candidate genes locate distantly while in a corrected assembly are next to each other. (b) Bar graphs show candidate gene specific allelic expression patterns. The blue bars represent the *X. maculatus* allele, and gray bars represent the *X. couchianus* allele. (c) The upper panel and lower panel shows allelic expression in *Sd-* and *Sd+* hybrids respectively. Bar heights represent allelic expression levels as in (d), and bars are plotted in the order of *X. maculatus* gene orders on Chr 13. Asterisks hallmark positions of candidate gene models. For alleles that are not expressed, they were assigned a 0 value for (b) and (d). (e) Dot plots show relative allelic expression of Chr 13 genes in *Dr+ Sd-* and *Dr+ Sd+* hybrids. For each dot, the X-coordinate represent relative expression levels of *X. maculatus* over *X. couchianus* alleles in *Dr+ Sd-* hybrids, and the Y-coordinate represent relative expression levels of *X. maculatus* over *X. couchianus* alleles in *Dr+ Sd+* hybrids. The dashed lines that located at Log_2_0.1 and Log_2_10 that correspond to relative expression ratio of 0.1 or 10 separates the graphs in 9 regions. Dots in the top left and bottom right regions represent genes that dominantly expressed for one allele in *Sd-* hybrids but the other allele in *Sd+* hybrids. Light gray dots are all genes on the Chr 13, with dark gray dots highlighted all genes in the Figure 2 Manhattan plot shoulder region. Red dots highlighted genes exhibit reversed dominant allelic expression between *Sd-* and *Sd+* hybrids. The dark red spot highlighted *long neurotoxin OH57* like. Allelic gene expression patterns for Chr 21 and Chr 13, and genes underlying *Sd* and *Dr* candidate regions are shown.

All *Sd +* hybrids exhibit *X. maculatus* allele specific expression for *long neurotoxin OH-57* and *sperm acrosome membrane-associated protein* (Fig 4c,d; Supplement Figure S3), but *X. couchianus* allele dominant expression for *long neurotoxin OH-57*, or bi-allelic expression for *sperm acrosome membrane-associated protein* in hybrids that do not exhibit *Sd* (Fig 4c,d). In addition, we surveyed the allelic expression patterns of all Chr 13 genes, *long neurotoxin OH-57*, and two additional genes (i.e., *fatty acid-binding protein 9-like* and *gastrula zinc finger protein XlCGF57.1-like*) that located on 6.38 Mbp and 6.63 Mbp regions are the only three genes exhibiting a *X. couchianus* allele dominant expression in *Dr+ Sd-*, and *X. maculatus* allele dominant expression in *Dr+ Sd+* hybrids [*X. maculatus/X. couchianus*<0.1 in *Dr+Sd-* hybrids and *X. maculatus/X. couchianus* >10 in *Dr+Sd+* hybrids, p-value < 0.05; Fig 4e].

The *long neurotoxin OH-57* gene is the only gene that exhibited a lower expression in *Dr+ Sd+* hybrids (Table 1), a genotype distribution bias from Hardy Weinberg Equilibrium (HWE), and reversed allelic expression patterns between *Dr+ Sd-* and *Dr+ Sd+* hybrids. The *X. maculatus* and *X. couchianus long neurotoxin OH-57* alleles exhibit two amino acid changes, A120T and G122S (Fig 5a). These two amino acids differences led to an shortened alpha-helix for the *X. couchianus* allele protein product and substitute a hydrophobic alanine to hydrophilic threonine (Fig 5b).

**Figure 5.**
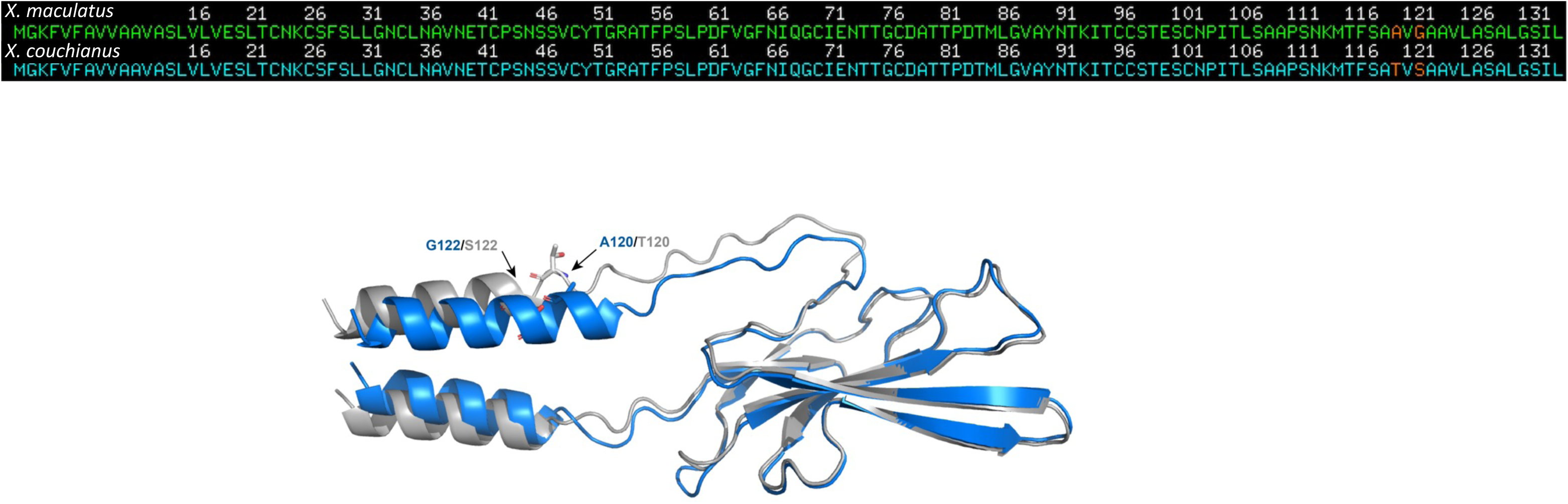
Structural comparison between *X. maculatus* and *X. couchianus* alleles for long neurotoxin OH-57-like peptide. (a) Primary peptide sequences comparison between the *X. maculatus* and *X. couchianus* alleles of long neurotoxin OH-57-like exhibit two amino acid changes (b) Structural alignments of long neurotoxin OH-57-like peptides derived from *X. maculatus* (blue) and *X.couchianus* (grey) using the align function of PyMOL.

### Macromelanophore can be modulated by epinephrine regulation

Melanosome trafficking is mediated by cAMP concentration (low cAMP triggers melanosome aggregation and high cAMP triggers melanosome dispersion). It was showed that melanosomes aggregate following epinephrine exposure in *Gambusia holbrooki* melanophores. Since we showed that a *long neurotoxin OH-57* is involved in the macromelanophore patterning, we hypothesized that *Xiphophorus* macromelanophore is subjected to neuronal regulation. We treated *Xiphophorus* fish with macromelanophores patterns, including *X. xiphidium* and F_1_ hybrid between *X. maculatus* and *X. couchianus* with 2.4 mg/ml epinephrine. Macromelanophores exhibited melanosome aggregation following 5min and 10min of epinephrine treatment in *X. xiphidium* (Fig. 6a) and *X. maculatus-X. couchianus* hybrids (Fig. 6b) respectively.

**Figure 6.**
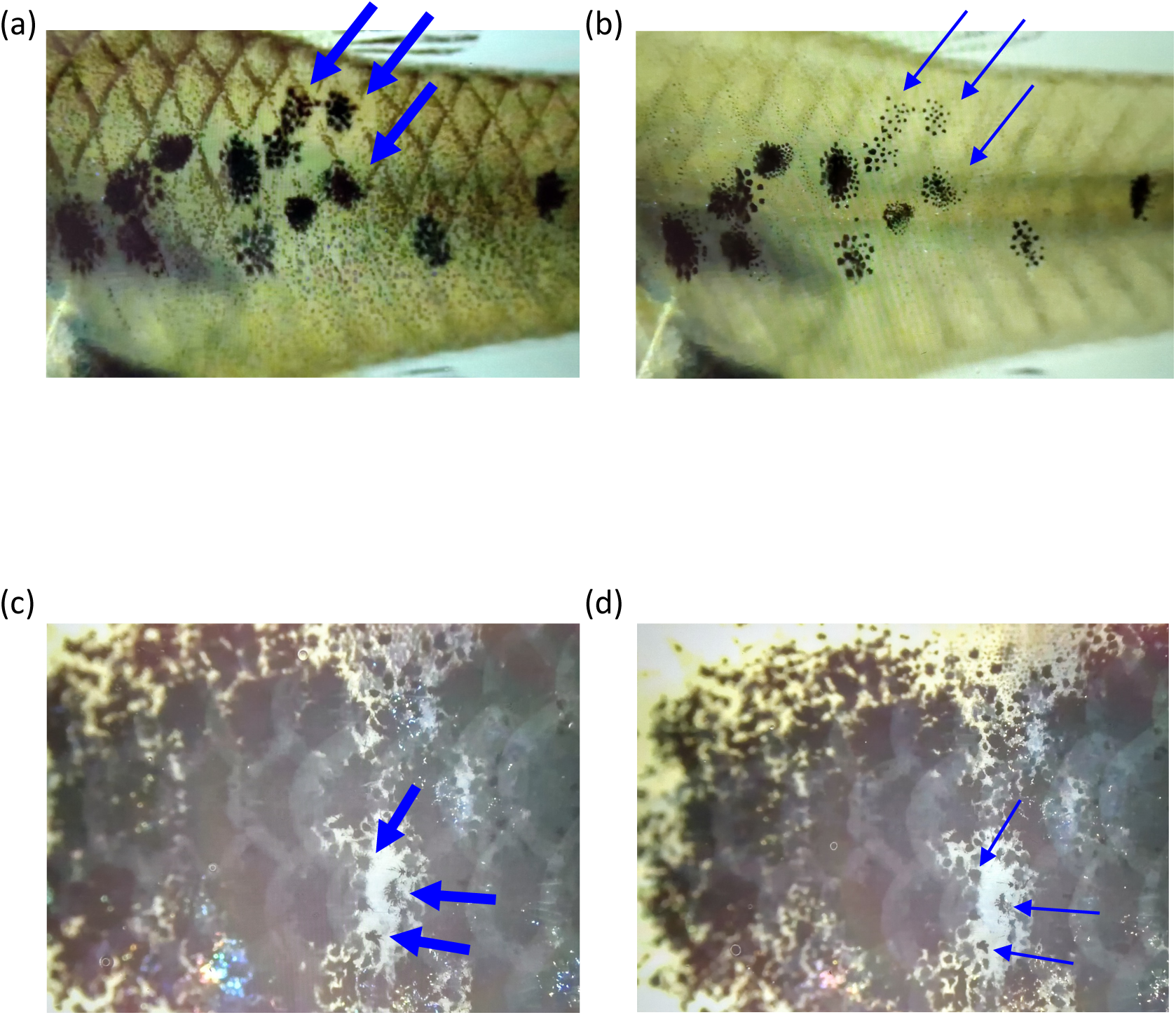
Epinephrine treatment triggers melanosome aggregation in macromelanophore. Epinephrine (24mg/ml) were treated to *X. xiphidium* (a-b), and interspecies hybrid between *X. maculatus* and *X. couchianus* (c-d). (a) Before treatment. (b) 5-min after treatment. (c) Before treatment. (d) 10-min after treatment.

## Discussion

In this study, we identified the long-sought loci underlying an ornamental trait *Dorsal red* (*Dr*) and disease related trait, *Spotted dorsal* (*Sd*) in *Xiphophorus* fish (21).

Our finding showed the *Dr* is a *X. maculatus* monogenic dominant trait with 100% genetic penetrance. This is well supported by our breeding records of hybrids involving *X. maculatus*, which display *Dr*, and from observations that phenotypical distribution of *Dr* follows a pattern that is consistent with expectations from a single dominant locus (Supplemental Table S1; Supplemental Table S2; Fig. 2a). It is important to note that *Dr* is a xanthophore pigmentation pattern and should be distinguished from the *xmrk* gene. The *xmrk* gene is a driver for cellular proliferation (22–26), while the *Dr* locus serves compartmentalization function that determines the spatial expression pattern of *xmrk*. In this regard, the cadherin genes (*cdh6* and *cdh20*) within the *Dr* locus are strong candidate genes accounting for the *Dr* phenotype.

Second, the loci underlying *Sd* and *Dr* are proximate on the *X. maculatus* sex chromosome Chr 21 but are regulated differently. The *Sd* and *Dr* pigmentation patterns are always linked in hybrids between *X. maculatus* and *X. hellerii* (13, 27), and were thus thought to be regulated simultaneously. In this study, *Dr* and *Sd* are mapped to the same region on Chr 21. This observation supports the initial thought that *Sd* and *Dr* are physically linked. However, unexpectedly, we showed that despite loci underlying *Dr* and *Sd* being proximate to each other, the two traits are not necessarily co-regulated, as shown in hybrids between *X. maculatus* and *X. couchianus* can display *Dr* independent of *Sd*. The finding that low expression of an autosomal locus from the *X. maculatus* (Chr 13) is needed for *Sd* expression suggests that *X. couchianus* allele of the locus can suppress *Sd*. We found that *long neurotoxin OH-57* gene is a candidate gene for regulating the *Sd*, evidenced by its *X. maculatus* allele-specific low expression in *Sd* + hybrids, and the expression of *X. couchianus* allele in *Sd -* hybrids. However, the suppression of *Sd* is likely mediated by more than the *long neurotoxin OH-57* locus, as *Sd* is observed in only 10% of the F_2_ hybrids, which is lower than an otherwise expected 25% for two-loci interaction, and the presence of 6 *Dr+ Sd-* hybrids that showed lowly expressed *X. maculatus* allele-only expression for this gene. The *Sd* phenotype exhibition is known to be varied in age. Even isogenic population can display this phenotype at different ages. Since all F_2_ hybrids were euthanized for sample collection at 9-month-old, the discrepancy in percentage of observed *Sd* + hybrid and anticipated *Sd* + hybrid from a two-gene model can also be sourced to a delayed development of *Sd*. However, from preliminary observations of aged F_2_ hybrid population (20 months-old and above), delayed *Sd* development has not been observed. Taken the *Sd* and *Dr* genetic mapping results together, we conclude *Sd* and *Dr* are two cell type-specific regulatory sequences for *xmrk*, with *Sd* for macromelanophore lineage, and *Dr* for xanthophore lineage (Fig. 7).

**Figure 7.**
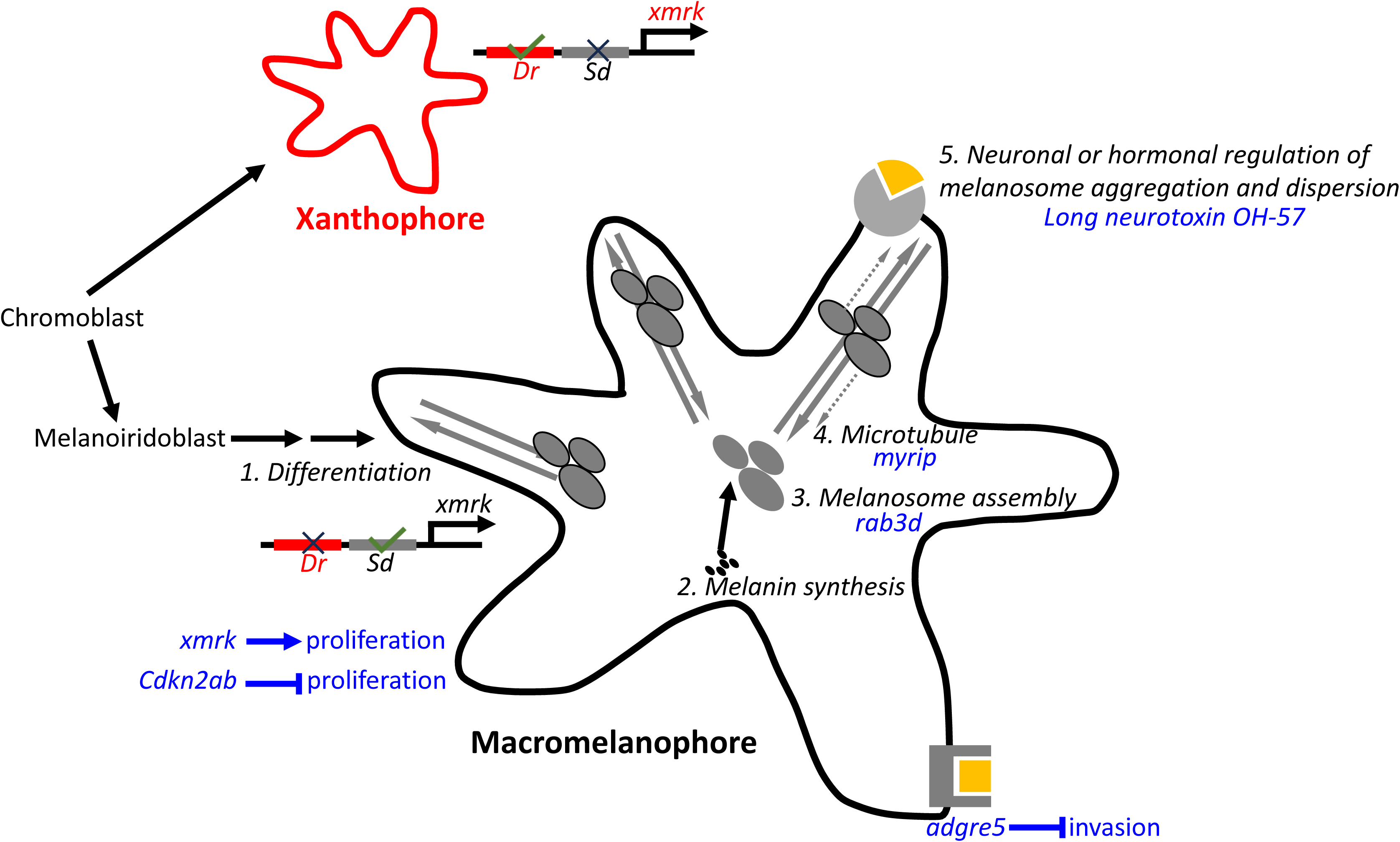
Overview of *xmrk* and regulator activity Table Legend.

The *Sd* pattern is known to be driven by *xmrk* oncogene. Genes suppressing the *xmrk* function have been found from several genetic mapping studies utilizing hybrids between *xmrk* positive and negative species. These regulators include *myrip* and *adgre5* in *X. malinche – X. birchmanni* hybrids, *cdkn2ab* and *rab3d* in *X. maculatus – X. hellerii* hybrids. In all cases, segregation between *xmrk* and the functional alleles of the regulators leads to lethal melanoma. Therefore, these regulators were interpretated as co-evolved with the *xmrk* oncogene to fine-tune the activity of it and suppress its oncogenic function. The negative epistasis between *xmrk* and non-functional alleles of these regulators in varied *Xiphophorus* hybrids serve as evidence supporting the Dobzhansky-Muller hybrid incompatibility model that states negative epistatic interactions occurring between genes that are adaptive in different species account for hybrid incompatibility and act as a post-zygotic barrier shielding gene flow between different species. However, in this study, *long neurotoxin OH-57-like* allele of *X. couchianus*, a species that does not have *xmrk*, fully “suppresses” the *xmrk-*driven *Sd* phenotype from a different species. The apparent “suppression” of *Sd* can be interpretated as either inhibition or lack of enhancement of *Sd* by *X. couchianus* allele than the *X. maculatus* allele. Macromelanophore pigmentation patterns, including *Sd*, is sexually selected [for review, see (28)], then presenting it at an adequate level is favored by sexual selection despite uncontrolled upregulation of *xmrk* is deleterious (i.e., melanoma). In this respect, the *X. maculatus* allele of *long neurotoxin OH-57* is a *xmrk* regulator that co-evolved with *xmrk*, by enhancing the *Sd* pigmentation. The *xmrk* originated from a local *egfr* gene duplication event. Multiple *Xiphophorus* species have it. A recent *Xiphophorus* phylogeny indicates that *X. couchianus* ancestral species lost the *xmrk* gene (29). When *xmrk* was lost, the *X. couchianus long neurotoxin OH-57-like* could degenerate due to no purification co-selection (Supplemental Figure 4).

Expansion and enhancement of macromelanophore pigmentation patterns in *Xiphophorus* are linked to the *xmrk* activity. A macromelanophore is characterized by its large size and is multinucleated (30). The *xmrk* gene and a hypothetical locus tightly linked to *xmrk* are thought to control the proliferation and compartmentalization of macromelanophore, respectively. For instance, a different laboratory strain *X. maculatus* (i.e., *X. maculatus* Jp163B) are fixed for *Spot Side* (*Sp*), a different allele of macromelanophore patterning locus as *Sd*, exhibits macromelanophore on the body side (27). In order for macromelanophore to be functional, several conditions are required: 1. Differentiation of macromelanophore from melanoblast; 2. Functional melanin synthesis pathway; 3. Assembly of melanosome; 4. Microtubule for melanosome transportation; 5. Intact signaling cascade regulating melanosome aggregation and dispersion within melanophore (Fig. 7). The *xmrk* drives macromelanophore proliferation (25, 26, 31). Several genes have been found to regulate *xmrk* oncogenicity using interspecies hybrids involving species encoding *xmrk*. These include genes that are thought to directly interact with melanosome assembly and transportation *rab3d* and *myrip* (13, 32, 33). Additional regulators include cell cycle regulator *cdkn2ab* (33–35), and cell adhesion, invasion related gene *adgre5* (32). The current finding that the *xmrk-*driven macromelanophore presentation is reliant on allele-specific expression of a neurotransmission regulator gene (i.e., *long neurotoxin OH-57-like*) suggests that the presentation of macromelanophore pattern has an additional point of regulation that is subjected to epistatic interactions. It has been shown that melanophores can be regulated by neuronal or hormonal ligands (30, 36). We have also shown that macromelanophore is regulated by adrenergic receptor signaling, despite the adrenaline-induced melanosome aggregation is not complete as observed in micromelanophore. From the understanding of how long neurotoxin Oh-57 work, i.e., binding to nicotinic acetylcholine receptor (nAChR) and inhibits acetylcholine from binding to nAChR (37), the *long neurotoxin OH-57-like* is likely to be a neurotransmitter receptor regulator in *Xiphophorus*. Our simulated *Xiphophorus* long neurotoxin OH-5-like peptide binding to nAChR is consistent with experimental solved neurotoxin-nAChR binding structure in terms of binding site and conformation (Supplemental Figure 5). Collectively, this newly identified *xmrk* regulatory point is likely due to incompatibility between macromelanophore and neurotransmitter regulator between different species.

In conclusion, we have identified a locus underlying a sex chromosome linked ornamental trait that is tightly linked to an oncogene in the *Xiphophorus*. Despite physical proximity, the ornamental trait loci and the oncogene are subjected to different genetic regulation by an autosomal regulator.

## Materials and Methods

### Animals

*X. maculatus, X. couchianus,* first generation (F_1_) interspecies hybrids between *X. maculatus* and *X. couchianus*, and intercross hybrids between F_1_ hybrids used in this study were supplied by the *Xiphophorus* Genetic Stock (http://www.xiphophorus.txstate.edu/). The F_1_ interspecies hybrids between *X. maculatus* Jp163A strain fish and *X. couchianus* were produced by enforced mating, and F_2_ hybrids were produced naturally by F_1_ hybrids. All fish were kept and samples taken in accordance with protocols approved by Texas State University IACUC (IACUC9048). All fish were dissected at age of 9-month. At dissection, all fish were anesthetized in an ice bath and upon loss of gill movement sacrificed by cranial resection. Dorsal fin clips were dissected and preserved in ethanol. Organs were dissected into RNAlater (Ambion Inc.) and kept at −80 °C until use.

### Phenotyping

Two pigmentation patterns, the macromelanophore pattern *spot dorsal*, and xanthophore pattern *dorsal red*, were typed. Before dissection, fish were nailed to a grading paper by using dissection needles to stabilize dorsal fin and anal fin. The *spot dorsal* is determined by a large black pigmented spot on the dorsal fin. This spotting pattern is different from the common micromelanophore pattern that is observed in all *Xiphophorus* fishes, which is smaller in size and distributed between fin rays on the dorsal fin. The *dorsal red* is determined by red coloration on the dorsal fin. The phenotyping results were agreed by 4 independent typing.

### DNA and RNA isolation

Fin clip of 4 *X. maculatus* males and 5 *X. couchianus* males were collected and digested by Protease K at room temperature for 1 hr. The lysate was then transferred to 2.0 ml collection tubes. DNA isolation was performed by a QIAcube HT (Qiagen) automated bio-sample isolation system, with reagent contained in QIAamp 96 DNA QIAcube HT Kit. The isolation system is equipped with a robotic arm with 8 pipettes. Each pipette is able to pick and eject pipette tips, self-clean, and transfer liquids between wells/columns, or between master reservoirs and wells/columns in standard 96-well plate formats. Each sample was independently maintained throughout the isolation process. Concentrations of DNA samples were measured using Qubit 2.0 fluorometer (Life Technologies, Grand Island, NY, USA), and adjusted for sequencing library preparation. Skin samples were homogenized in TRI-reagent (Sigma Inc., St. Louis, MO, USA) followed by addition of 200 μl/ml chloroform, vigorously shaken, and subjected to centrifugation at 12,000 g for 5min at 4°C. Total RNA was further purified using an RNeasy mini RNA isolation kit (Qiagen, Valencia, CA, USA). Column DNase digestion at 25°C for 15 min removed residual DNA. Total RNA concentration was determined using a Qubit 2.0 fluorometer (Life Technologies, Grand Island, NY, USA). RNA quality was verified on an Agilent 2100 Bioanalyzer (Agilent Technologies, Santa Clara, CA) to confirm that RIN scores were above 7.0 prior to subsequent gene expression profiling.

### Inter-specific genetic variants identification and annotation

To identify interspecies polymorphisms between the *X. maculatus* and *X. couchianus*, genomic DNAs of 4 *X. maculatus* and 5 *X. couchianus* were isolated and forwarded for genome shotgun sequencing library preparation using Illumina Nextera sequencing Library Prep Kit, followed by sequencing on HiSeq 2000 (Illumina, Inc., San Diego, CA, USA) using 150 bp paired-end (PE) sequencing strategy. Raw sequencing reads were filtered using fastx_toolkit (http://hannonlab.cshl.edu/fastx_toolkit/index.html). Filtered sequencing reads were mapped to the reference *X. maculatus* genome (GenBank assembly accession: GCA_002775205.2) using Bowtie2 “head-to-head” mode (38). Alignment files were sorted using Samtools (39). Pileup files were generated for each *X. maculatus*, and *X. couchianus* sample, and variant calling was processed by BCFtools for polymorphisms detection, with minimum variant locus coverage of 2 and variant genotyping call Phred score of 0 and alternative genotyping Phred score ≥ 20 for BCFtools (39–41). Genotype refers to inheritance of ancestral alleles, with heterozygous meaning that a locus exhibited genetic material from both ancestors (i.e., *X. maculatus* and *X. couchianus*), and homozygous means that a locus exhibited genetic material from only one parental species. Inter-specific polymorphic sites where all sequenced *X. maculatus* support a homozygous *X. maculatus* reference, and *X. couchianus* support a homozygous alternative variant calls were kept as a reference of genetic variance in .bed format.

### Gene expression profiling and allelic expression profiling

RNA sequencing was performed upon libraries constructed using the NEB stranded mRNA-seq library preparation kit (New England Biolabs, Ipswich, MA, USA). RNA libraries were sequenced as 150bp pair-end fragments using Illumina Novaseq system (Illumina, Inc., San Diego, CA, USA). RNA libraries were sequenced as 150bp paired-end fragments using Illumina NovaSeq system (Illumina, Inc., San Diego, CA, USA). Short sequencing reads were filtered using fastx_toolkit (http://hannonlab.cshl.edu/fastx_toolkit/contact.html). RNA-Seq sequencing reads were produced from independent skin samples of F_2_ hybrids exhibiting *Sd* and only *Dr*. Sequencing reads were mapped to *X. maculatus* reference genome (GenBank assembly accession: GCA_002775205.2) using Tophat2 (42, 43). Gene expression was subsequently profiled by counting number of sequencing reads that mapped to gene models annotated by NCBI using Subread package FeatureCount function (44). Differentially expressed genes were identified using R package edgeR, with p-value adjusted using False Discovery Rate (FDR) method (45). |Log_2_FC|>1 and FDR<0.05 was used to determine differentially expressed genes.

We used an established method to assess allelic expression of the F_2_ hybrids (46). Briefly, orthologs between the parental alleles were identified using blastn, with the *X. maculatus* transcripts as subject and *X. couchianus* as reference. Both parental alleles were subsequently combined to represent a hybrid transcriptome. RNA-Seq reads were mapped to this hybrid transcriptome using Bowtie2 (38), followed by filtering of the read mapping file to keep reads that were mapped uniquely to one allele, and quantification of these reads per allele. Both parental alleles read counts of a gene was normalized to the length of parental allele lengths and were used to estimate percentages of gene expression contributed by either parental allele. These ratios were subsequently used to calculate allele expression using the library size normalized total read counts. For data visualization, gene expression read counts were normalized to library size and were plotted as bar graph using custom scripts in R (v3.5.1).

### Genome mapping of *Spot Dorsal* and *Dorsal Red*

Sequencing adaptor contamination of RNAseq reads was first removed from raw sequencing reads using fastx_toolkit, followed by trimming of low-quality sections of each sequencing read. Low quality sequencing reads were further removed from sequencing result (http://hannonlab.cshl.edu/fastx_toolkit/index.html). Processed sequencing reads were mapped to *X. maculatus* genome v5.0 (GenBank assembly accession: GCA_002775205.2) using Bowtie2 (38). Mpileup files were made using legend version of samtools (v0.19) and genotyping was processed using Bcftools. Genotype in this study refers to inheritance of ancestral alleles, with heterozygous meaning that a locus exhibited genetic material from both ancestors (i.e., *X. maculatus* and *X. couchianus*), and homozygous means that a locus exhibited genetic material from only the one parental species. This samtools version performs Chisq test for genotyping distribution against Hardy Weinberg Equilibrium between two groups. Hybrid individuals that do not exhibit phenotype-to-test were assigned as group 0 and individuals that exhibit a phenotype were assigned as group 1. The Chisq test p-values were adjusted using Bonferroni method, and were converted to −10*Log_10_p-value for Manhattan plot. To assess individual genotype at reference inter-specific polymorphic sites, genotype calls were required to be supported by specific statistics (i.e., Bcftools: MAPQ >=30, Phred score of genotype call = 0, with alternative genotype call Phred score >=20).

### Protein structural simulation

The primary amino acid sequences of long neurotoxin OH-57-like peptide for *X. maculatus* (XP_023200269.1) and *X. couchianus* (XP_027891100.1) were retrieved from NCBI. The structural models of long neurotoxin OH-57-like peptides were obtained from the protein structural prediction software, ColabFold (38). ColabFold combines the homology search of Many-against-Many sequence searching (MMseqs2) (47) with AlphaFold2 (48) to predict protein structures and complexes. For each peptide, 5 models were generated and the model with the highest mean pLDDT score - 76.72 for *X. couchianus* and 76.80 for *X. maculatus* - was selected for alignment. The chosen models were then aligned and visualized using PyMOL, focusing on structural variations.

### Protein Docking and Visualization

The initial structure of the nicotinic acetylcholine receptor (nAChR) was obtained from the Protein Data Bank (PDB), specific as 2BG9. Only the extracellular region of the nAChR was used in the following docking process. The docking study to investigate the interaction between the long neurotoxin OH-57-like peptide, which obtained from Colabfold, and the extracellular region of nAChR was conducted using the ZDOCK (https://zdock.wenglab.org) and ZRANK programs. Initially, ZDOCK was employed to predict potential binding poses of the peptide to the receptor. ZDOCK utilizes a fast Fourier transform (FFT) algorithm to perform a comprehensive search of the rotational and translational space between the receptor and ligand, generating a set of possible docked conformations (49).1 Following the initial docking, the top-ranked poses from ZDOCK were further refined using ZRANK. ZRANK re-ranks the docking poses based on a detailed scoring function that considers additional energy terms, including van der Waals, electrostatics, and desolvation contributions (50). This refinement step ensures a more accurate prediction of the binding affinity and interaction specifics. The docking structure was visualized using PyMOL.

### Epinephrin treatment to *Xiphophorus*

Two types of *Xiphophorus* fish exhibiting macromelanophore pigmentation patterns, *X. xiphidium* (*RP* line) and F_1_ interspecies hybrid between *X. maculatus* (Jp163B) and *X. couchianus*, were provided by the *Xiphophorus* Genetic Stock Center. Fish were euthanized by over-dosing MS222. Following loss of gill movement, fish were decapitalized and immersed in 24mg/ml Epinephrin (Fisher Scientific, Hampton, NH). Macromelanophores were videotaped for 10 minutes.

### Inter-species chromosomal orthologous region identification

To identify genome region of *X. couchianus* that is orthologous to *X. maculatus* region underlying candidate *Dr* locus, we used Liftoff (https://github.com/agshumate/Liftoff) to map *X. maculatus* genes onto *X. couchianus*. After synteny filtering, we found genes within a ∼200kb area of *X. maculatus* chr21 failed to mapped onto *X. couchianus*, including *xmrk*. To further confirm this synteny gap, we used minimap2 (51, 52) to align the whole genome between the two species and improved the alignment with Genome Alignment Tools from the Hiller lab (53, 54).

## Acknowledgments

This work was supported by the National Institutes of Health, National Cancer Institute, R15 CA-223964 and R24 OD-031467 from the NIH Division of Comparative Medicine, R2R1 accelerator award from Texas State University. RNA-seq data was generated in the Genome Sequencing Facility, which is supported by UT Health San Antonio, NIH-NCI P30 CA054174 (Cancer Center at UT Health San Antonio) and NIH Shared Instrument grant S10OD030311 (S10 grant to NovaSeq 6000 System), and CPRIT Core Facility Award (RP220662), and NIH NCI R50 CA265339 to Z. Lai.

