## Supplemental Figure and Tables for "Segregation between an ornamental and a disease driver gene provides insights into pigment cell regulation": Supplemental FigureForArch.pptx

### Slide 1
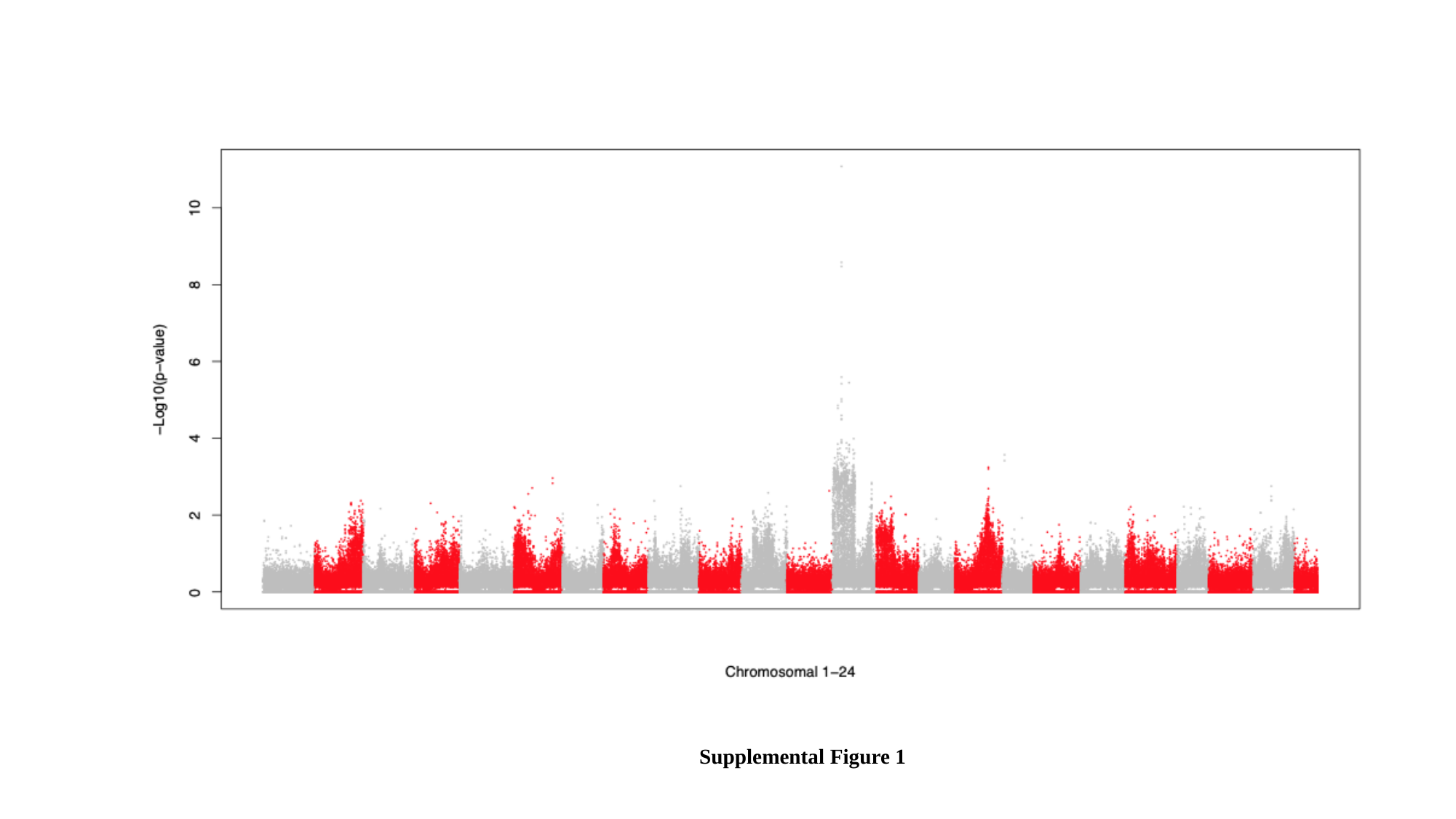

Supplemental Figure 1

### Slide 2
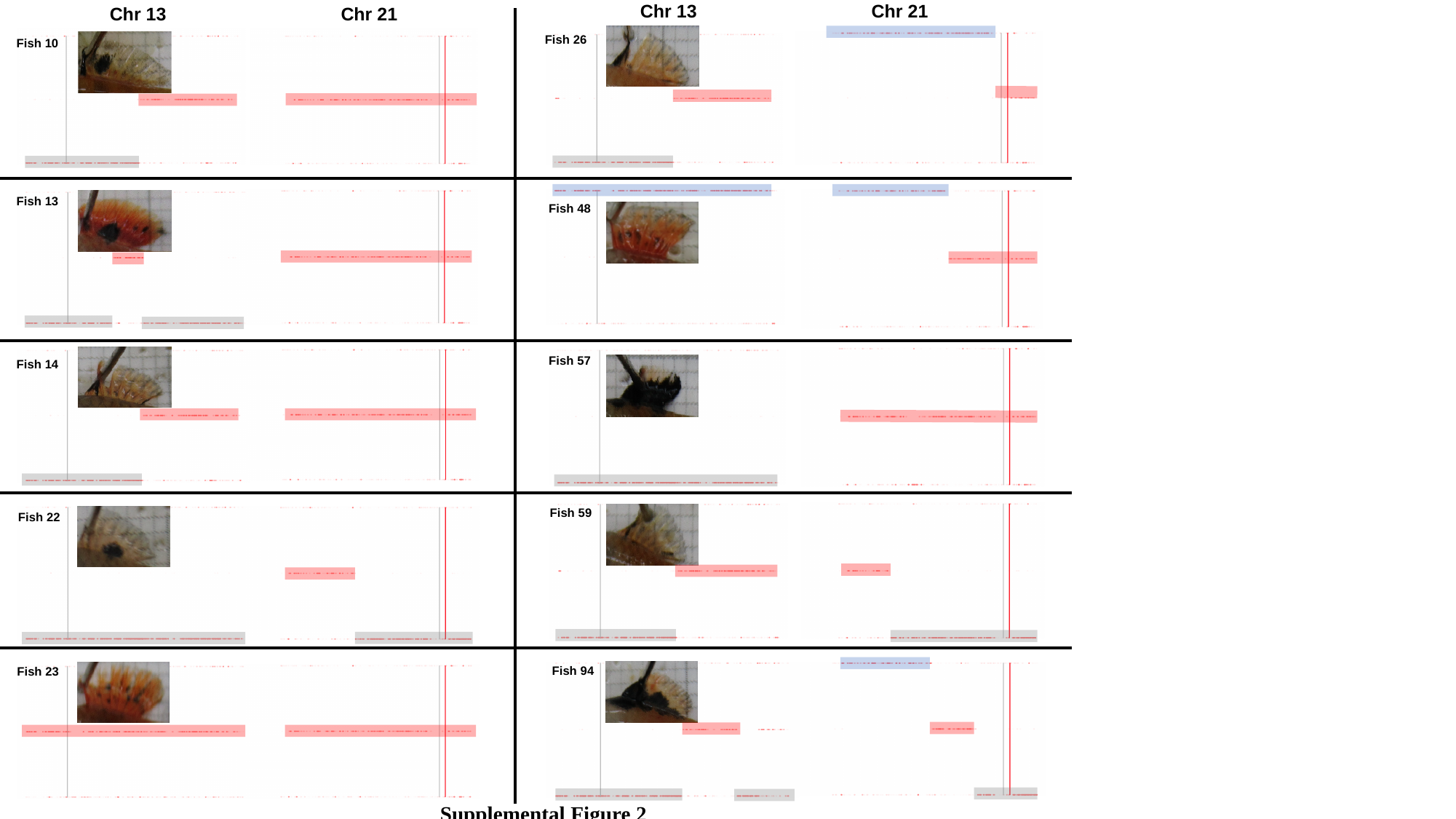

Chr 13
Chr 21
Chr 13
Chr 21
Fish 26
Fish 10
Fish 13
Fish 48
Fish 57
Fish 14
Fish 59
Fish 22
Fish 94
Fish 23
Supplemental Figure 2

### Slide 3
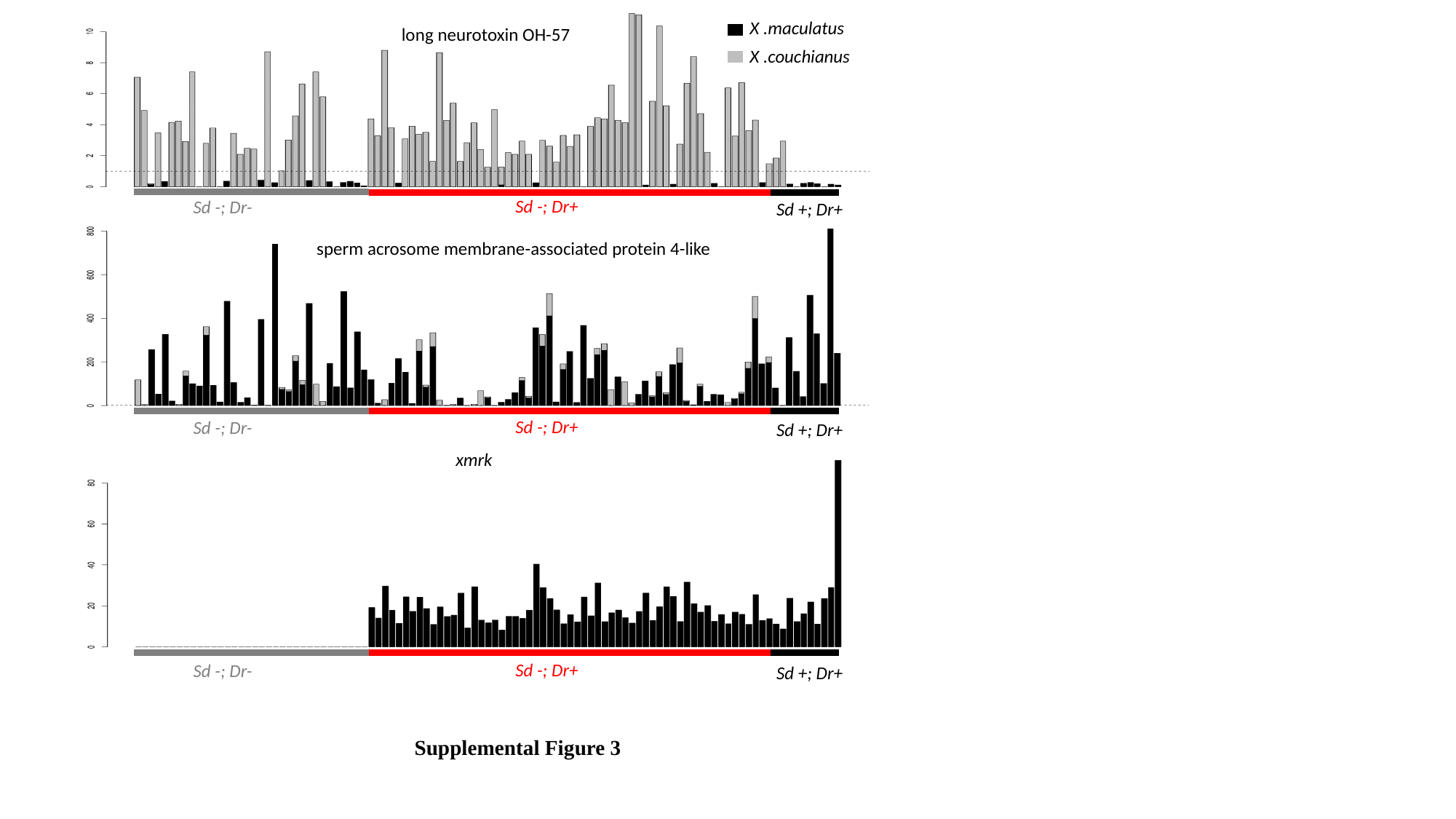

X .maculatus
long neurotoxin OH-57
X .couchianus
Sd -; Dr+
Sd -; Dr-
Sd +; Dr+
sperm acrosome membrane-associated protein 4-like
Sd -; Dr+
Sd -; Dr-
Sd +; Dr+
xmrk
Sd -; Dr+
Sd -; Dr-
Sd +; Dr+
Supplemental Figure 3

### Slide 4
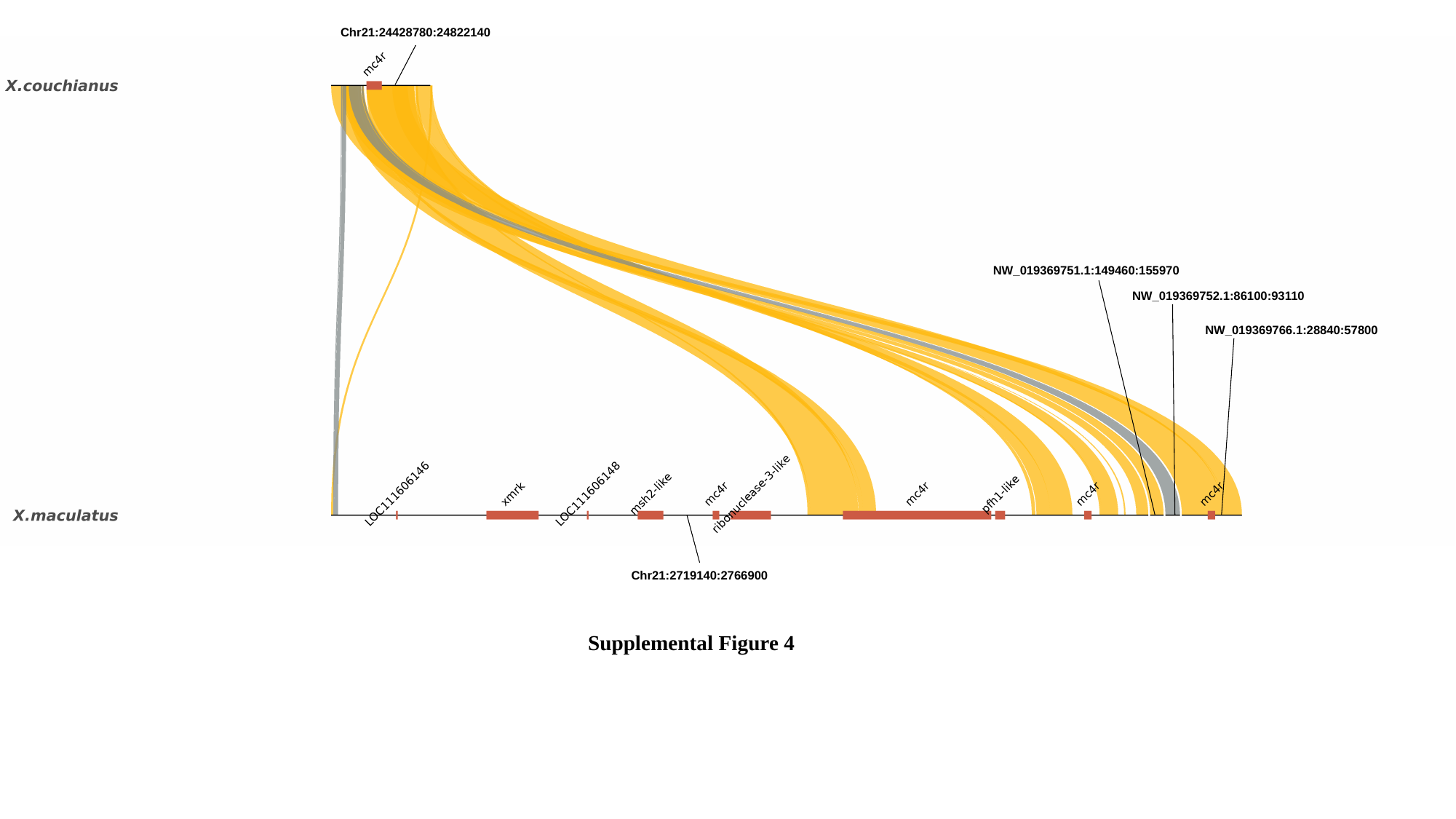

Chr21:24428780:24822140
NW_019369751.1:149460:155970
NW_019369752.1:86100:93110
NW_019369766.1:28840:57800
Chr21:2719140:2766900
Supplemental Figure 4

### Slide 5
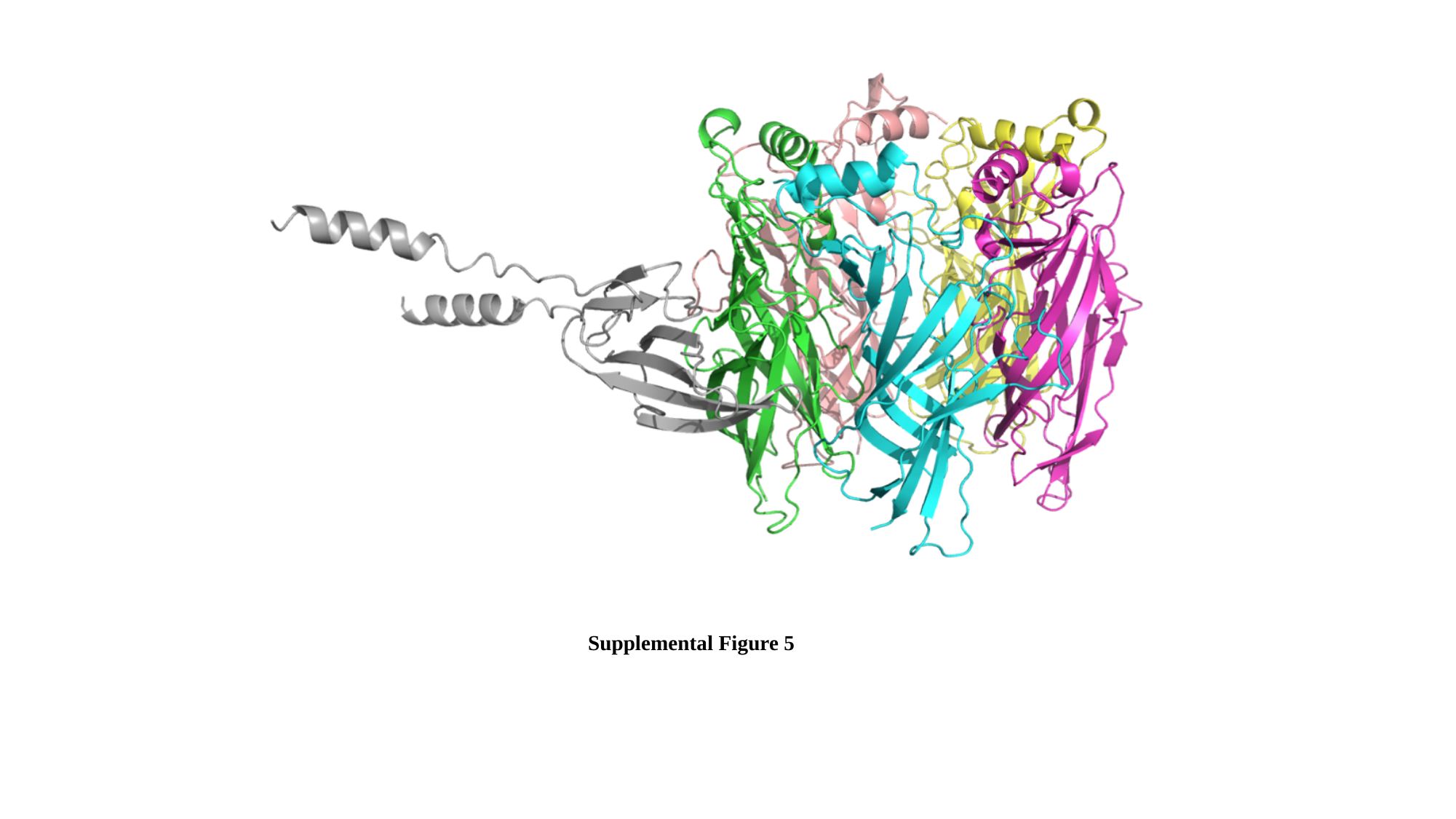

Supplemental Figure 5
